# When genomes collide: multiple modes of germline misregulation in a dysgenic syndrome of *Drosophila virilis*

**DOI:** 10.1101/005124

**Authors:** Mauricio A. Galdos, Alexandra A. Erwin, Michelle L. Wickersheim, Chris C. Harrison, Kendra D. Marr, Justin P. Blumenstiel

## Abstract

In sexually reproducing species the union of gametes that are not closely related can result in genomic incompatibility. Hybrid dysgenic syndromes represent a form of genomic incompatibility that can arise when transposable element (TE) abundance differs between two parents. When TEs lacking in the female parent are transmitted paternally, a lack of corresponding silencing small RNAs (piRNAs) transmitted through the female germline can lead to TE mobilization in progeny. The epigenetic nature of this phenomenon is demonstrated by the fact that genetically identical females of the reciprocal cross are normal. Here we show that in the hybrid dysgenic syndrome of *Drosophila virilis*, an excess of paternally inherited TE families leads not only to increased expression of these TEs, but also coincides with derepression of TEs in equal abundance within parents. Moreover, TE derepression is stable as flies age and associated with piRNA biogenesis defects for only some TEs. At the same time, TE activation is associated with a genome wide shift in the distribution of endogenous gene expression and an increase in abundance of off-target genic piRNAs. To identify regions of the maternal genome that most protect against dysgenesis, we performed an F3 backcross analysis. We find that pericentric regions play a dominant role in maternal protection. This F3 backcross approach additionally allowed us to clarify the properties of genic paramutation in *D. virilis*. Overall, results support a model in which early germline events in dysgenesis establish a chronic, stable state of mis-expression that is maintained through adulthood.

Such early events in the germline that are mediated by parent-of-origin effects may be important in determining patterns of gene expression in natural populations.

**Author Summary:** Transposable elements (TE) are selfish elements that code for the function of copying themselves. More than half the human genome is comprised of such elements. Studies in the fruit flies *Drosophila melanogaster* and *D. virilis* have been important in demonstrating a role for RNA silencing by piwi-interacting RNAs (piRNAs) in protecting the genome against these harmful elements. These small RNAs are capable of recognizing TE mRNAs and mediating their destruction by Argonaute proteins. They are also transmitted by the female germline to offspring in order to maintain a stable genome across generations. When males carrying a particular TE family are crossed with females lacking the element, the mother is unable to provide genome defense via complementary piRNAs that target the element. This leads to excess TE activation in the germline and sterility. This phenomenon is known as hybrid dysgenesis. In this article we characterize the genomic landscape of TE destabilization that occurs in hybrid dysgenesis in *D. virilis*. Previous studies had demonstrated that multiple TEs mobilized during hybrid dysgenesis. We demonstrate that this mobilization of multiple TEs is associated with activation of additional TEs in the germline. In addition, we find that TE activation leads to the production of off-target genic piRNAs that cause reduced expression of highly expressed genes. Finally, we show that genic off-target effects of piRNA silencing can contribute to parent-of-origin effects on gene expression. Similar phenomena may influence patterns of gene expression in the germline of natural populations.

## Introduction

In sexually reproducing species, two unique haploid genomes join together in syngamy to establish each generation. This mixing of genomes introduces potentially advantageous variation under changing environmental conditions, but also provides a condition ripe for exploitation by selfish elements [1]. Because syngamy can introduce selfish elements to new genomes and recombination can separate selfish elements from their harmful consequences, selfish elements such as transposable elements (TEs) can proliferate [2,3]. When the balance of TE burden differs between gametes of mated individuals, a genomic clash known as hybrid dysgenesis may occur [4,5]. Hybrid dysgenesis is associated with sterility and an increased mutation rate that typically occurs when males with a particular TE family are mated with females lacking that family [6,7,8]. Sterility and mutation occurs because paternally inherited TEs become over-activated in the F1 germline [9]. In *Drosophila melanogaster*, three distinct hybrid dysgenic syndromes have been identified. These are driven by the P element, the I element and the hobo element [6,7,8]. In addition, a second form of hybrid dysgenesis has been identified in *D. virilis* [10]. The *D. virilis* syndrome has been proposed to be driven by the *Penelope* element [11,12,13,14], though other elements may contribute to dysgenesis [15,16].

Hybrid dysgenesis syndromes in *Drosophila* have provided crucial insight into TE dynamics and mechanisms of host genome defense by small RNAs. The classical observation – activation for TEs inherited through the *Drosophila* male germline – can be explained by the fact that the maternal germline is the primary agent of transgenerational TE repression via small RNA silencing pathways [17]. Mothers lacking a particular TE family in their genome also lack corresponding 23-30 nt piwi-interacting RNAs (piRNAs) in their eggs. In the absence of the maternally provisioned piRNAs that target TE mRNAs for Argonaute-mediated slicing, paternally inherited TEs become activated in the progeny germline. This has been demonstrated for the P element and the I element systems of hybrid dysgenesis in *Drosophila melanogaster* [18,19,20,21].

It remains unclear what precisely follows the initial germline activation of paternally inherited TEs. Early studies of P element hybrid dysgenesis in *D. melanogaster* indicated downstream activation of additional TEs [22], but this interpretation was soon called into doubt [23]. Nonetheless, the syndrome of hybrid dysgensis in *D. virilis* provided strong support for cascading germline activation of TEs because multiple TEs were found to mobilize in the germline of dysgenic progeny [11,15]. While different TE families may contribute to the initial induction of dysgenesis in *D. virilis*, germline co-mobilization is demonstrated for two TEs (*Ulysses* and *Telemac*) that are evenly distributed between the two strains and also mobilize during dysgenesis [16,24,25]. Significantly, a recent study of the P element system has indicated that the previous conclusion of no co-mobilization may have been premature [21]. In the face of P element activation, DNA damage can perturb piRNA biogenesis and this defective piRNA biogenesis in turn leads to mobilization of TEs. Thus, global TE mobilization may be common in syndromes of hybrid dysgenesis that are driven by a single element. Whether the mechanism underlying global TE activation is shared across syndromes is not known.

Here we use the unique system of hybrid dysgenesis in *D. virilis* to define the landscape of TE derepression and genic mis-expression that is driven by the initial activation of a small number of TE families. We show that germline mobilization of TEs is driven by a multi-layered mechanism. Diverse elements are activated corresponding to TE copy number asymmetry between strains and there is a corresponding activation of multiple TEs that are evenly distributed between strains. This state of chronic TE derepression is maintained as flies age. In turn, highly expressed genes become *de novo* targets of piRNA biogenesis, which leads to a shift in the balance of endogenous gene expression.

## Results

### Genome Wide Asymmetry in TE abundance in a Dysgenic Cross of *D. virilis*

Previous studies of hybrid dysgenesis in *D. virilis* have identified several candidate elements that appear to contribute to F1 sterility in crosses between inducer strain 160 fathers and the non-inducer strain 9 mothers. The *Penelope* element is most likely the dominant element. Active copies of *Penelope* are abundant in strain 160 and only degenerate copies are present in strain 9 [26]. Furthermore, expression of the *Penelope* element is elevated in the ovaries and testes of F1 dysgenic progeny that have escaped ablation of the gonad [11]. In addition to the *Penelope* element, three other elements (*Helena, Paris* and *Polyphemus* [16,27]) are also more abundant in the inducer strain, and these likely contribute to dysgenesis. A complex mode of hybrid dysgenesis, driven jointly by multiple elements, is supported by the fact that some strains of *D. virilis* lack *Penelope* piRNAs in the ovary but are “neutral” - capable of preventing dysgenesis when crossed with strain 160 males but also incapable of induction [28]. If *Penelope* is the sole cause of paternal induction, it is difficult to explain how these strains could prevent induction when the female germline lacks *Penelope* piRNAs.

To determine whether additional elements to *Penelope* may contribute to dysgenesis in *D. virilis*, we performed genome sequencing on both strains 9 (non-inducer) and 160 (inducer). To estimate bulk differences in TE abundance between the two strains, we mapped single end reads against a custom *D. virilis* TE library (Supplemental 1) that included more TEs than used in the previous analysis of *Polyphemus* [27]. Relative mapping abundance with BWA-MEM[29], normalized by read number mapping to the reference, is shown in Figure 1A. Consistent with previous results, *Penelope* and *Polyphemus* show the greatest excess in strain 160. Using a 3-fold cutoff as a threshold, we also confirm the previously reported result that copy number for *Helena* and *Paris* is enriched in strain 160[16]. Furthermore, we identify seven additional elements that are greater than 3-fold enriched in strain 160. These elements are all candidates for contributing to the dysgenic syndrome. Interestingly, the telomeric TART elements exhibit higher mapping abundance in strain 160, albeit below the 3-fold enrichment threshold.

**Figure 1.**
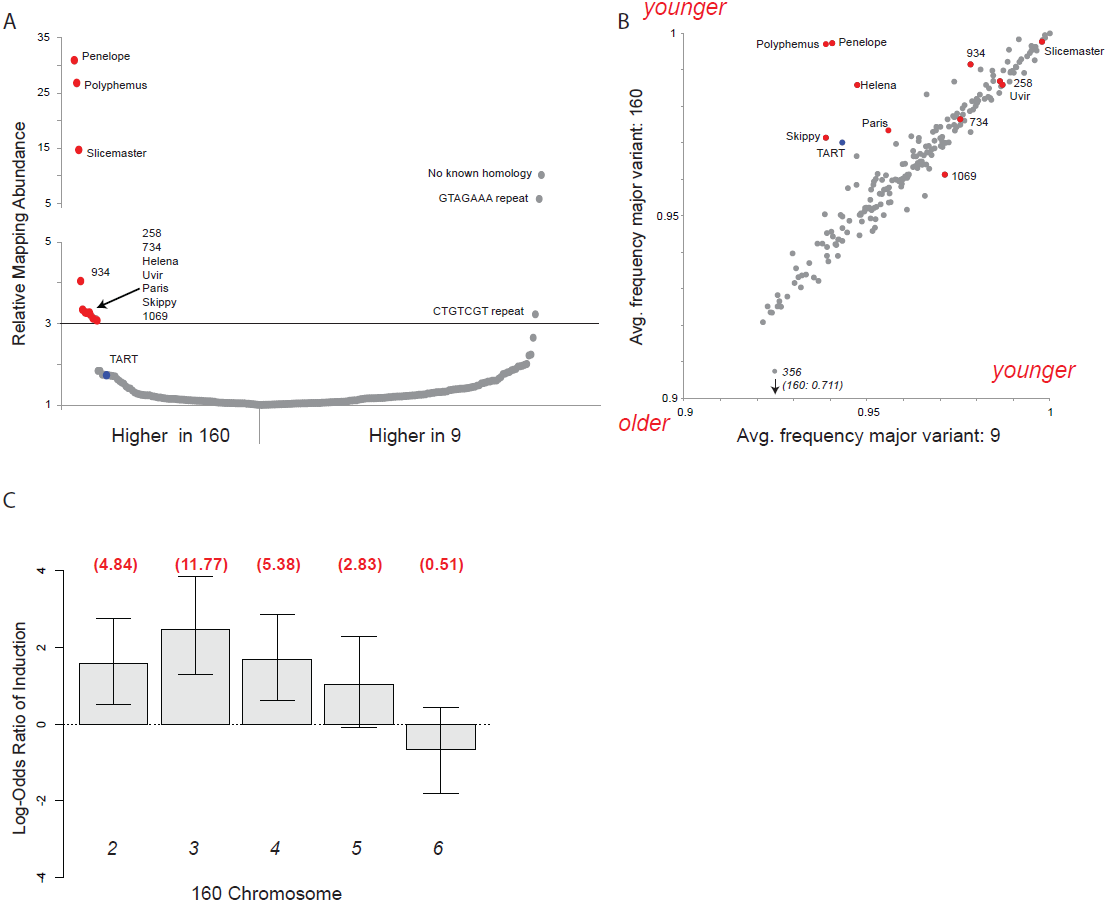
Multiple transposable elements are associated with induction of hybrid dysgenesis. (A) Relative mapping abundance of single-end, 100 bp reads from strain 9 and strain 160 (normalized by reads mapping to the genome), to a consolidated repeat library. Eleven elements are in 3-fold excess in strain 160. TART elements are about 1.7-fold in excess. No apparent TEs were found in excess in strain 9. (B) Using piledriver (https://github.com/arq5x/piledriver) we assessed homogeneity within reads mapping to the TE library by determining the average frequency of the major variant in both strains. TEs in excess in strain 160 are either more homogenous in strain 160, or similarly aged between strains. (C) Induction of sterility by 160 is broadly distributed across the genome, with the exception of chromosome 6 (the dot chromosome). Log odds ratios for probability of induction were estimated by crossing F1 males to strain 9, determining whether F2s had male gonadal atrophy and genotyping F2s to determine the chromosomes inherited from the father. Estimates were determined using a generalized linear model for logistic regression (binomial family with a logit link). Values in red are actual odds ratios. Whiskers are 95% confidence intervals. Chromosome 5 is significant at 0.1 level only. X chromosome is not scored because dysgenesis is scored in males and males do not inherit the X from their fathers (N=92).

Thus, it appears that diverse TE families are in excess in strain 160. This is consistent with the hypothesis that the invasion of the *Penelope* element itself has contributed to genome instability by perhaps mobilizing other TEs within the strain[13]. By contrast, elements enriched in strain 9 do not appear to be transposable elements. However, it is important to note that ascertainment bias may contribute to the observation that no TEs appear in significant excess in strain 9. This is because the reference genome is more closely related to strain 160, and TEs that are entirely unique to strain 9 may be missing in the TE libraries generated through computational means from the reference genome.

### TE age analysis identifies different modes for TE asymmetry between strains

TE abundance between strains can result from different processes. For example, long-resident TEs may be in excess in one strain due to strain or population specific recent re-activation. By contrast, entirely new TEs may have invaded a species and have yet to spread equally throughout the genomes of the individuals within the population. The P element invasion in *D. melanogaster* is an example of the latter process [30]. It has recently invaded and is only present in current natural populations, not strains collected seven or more decades ago.

To distinguish among alternative processes contributing to asymmetry in TE abundance, one can perform an age analysis of given TE families. High sequence homogeneity within a TE family is an indicator of recent activation or invasion. A phylogenetic approach using full-length fragments is ideal for this purpose, but full-length TE assemblies are not available with short read sequencing technology. Therefore, we estimated TE family “age” by examining the sequence heterogeneity within mapping reads (Figure 1B). Age was estimated by considering the average frequency of the most common nucleotide, across nucleotides within the mapping. A young element that has recently invaded will show high similarity among copies, nearing 1 for an average frequency of the major nucleotide variant. Older elements, with patterns of activation that occurred more distantly in the past, accumulate differences among insertions. This accumulation of differences among multiple copies is made evident by lower nucleotide frequencies of the most common variant. Figure 1B plots these age measures for each TE within the two strains.

This analysis reveals two classes of TEs that are enriched in inducer strain 160. Consistent with previous analysis[26,27,31], *Penelope* and *Polyphemus* show greater homogeneity among copies in strain 160. This pattern also applies, albeit to a lesser extent, to *Helena*, *Paris, Skippy* and the telomeric TART element. This pattern indicates recent activation from long-term resident status.

A second class of elements with excess in strain 160 show a different pattern. *Slicemaster*, *Uvir, 258* and *734* are all either similarly aged within strains or, in the case of *1069*, slightly older in strain 160. In the case of elements like *Slicemaster*, which are very young (>99% similarity), this can be explained by recent invasion of both strains but excess movement in strain 160, rather than “re-activation” from long-time resident status.

### The genomic distribution of paternal induction in hybrid dysgenesis

Previous studies have indicated that the *Penelope*, *Paris* and *Helena* elements are broadly distributed across multiple chromosomes of the inducer strain 160 [16]. Considering the much larger number of TEs that are now known to be in excess in strain 160, we sought to determine the strength of induction across paternally inherited 160 chromosomes. This was achieved by generating F1 males heterozygous for strain 160 and strain 9 chromosomes (through a non-dysgenic cross), crossing these males to strain 9 females, scoring for dysgenesis in F2 progeny, and genotyping F2s for chromosomes inherited from the heterozygous father. Overall, we found that induction of dysgenesis is distributed across all tested chromosomes, with the exception of the dot sixth chromosome (Figure 1C). Thus, an excess of diverse TEs in strain 160 is associated with induction distributed across the genome. In light of this, we sought to determine how TE excess in strain 160 corresponded with patterns of increased TE expression in progeny of the dysgenic cross.

### Patterns of TE expression in a dysgenic cross

To determine how TE excess in strain 160 corresponds with TE expression in dysgenic progeny, we performed mRNA-seq from pooled 0-2 hour old embryos laid by F1 females of both the dysgenic (9 females X 160 males) and non-dysgenic (160 females X 9 males) directions of the cross. One may also measure expression in F1 ovaries, but here this is not preferred since dysgenic ovaries are atrophied and expression analysis from these tissues is confounded by altered ratios of somatic and germline tissue. 0-2 hour old embryos laid by F1 mothers represent a sample of pure germline tissue since zygotic transcription in *D. virilis*, as measured with the early, zygotic *fushi-tarazu* (*ftz*) gene, begins after 2 hours [32]. Confirmation that embryos in this sample were collected prior to the onset of zygotic transcription was obtained by inspecting expression of *ftz*.

Full penetrance of dysgenesis, evidenced by atrophied gonads, is observed in approximately 50% of male and female progeny from 9 female X 160 male crosses. Therefore, embryos analyzed by mRNA-seq were those laid by mothers that escaped full sterility. For clarity, these tissues will be referred to as dysgenic, even though these tissues escaped complete atrophy. Sexual maturity in *D. virilis* is at about 5 days. To determine the dynamics of TE expression as flies aged, we analyzed mRNA-seq data from embryos laid by F1 mothers 12 to 16 days old, as well as 19 to 21 days old.

mRNA-seq analysis of dysgenic and non-dysgenic germline indicated different modes of TE mis-expression. TEs with excess abundance in the inducer strain were broadly overexpressed in the dysgenic germline. There were high magnitude differences for some elements such as *Skippy* and *Helena* but for others, such as *Paris*, *Polyphemus* and *TART,* differences were more modest (Figure 2A and B). Coupled with this, there was also a global pattern of overexpression for all TEs in the dysgenic germline, as evidenced by RPKM levels above the diagonal (Figure 2A and B). This pattern of global TE derepression applies largely to TEs that are lowly expressed. It should be noted that some caution must be taken in the interpretation of lowly expressed TEs. RPKM is fundamentally a proportional measure which is sensitive to changes in expression of highly expressed genes (see below). Nonetheless, visual inspection shows that there are at least three elements, not members of the class enriched in strain 160, that show increased expression in the dysgenic germline and are not lowly expressed (>8 RPKM). Importantly, expression level differences do not ameliorate with age of mother (compare Figures 2A and B).

**Figure 2.**
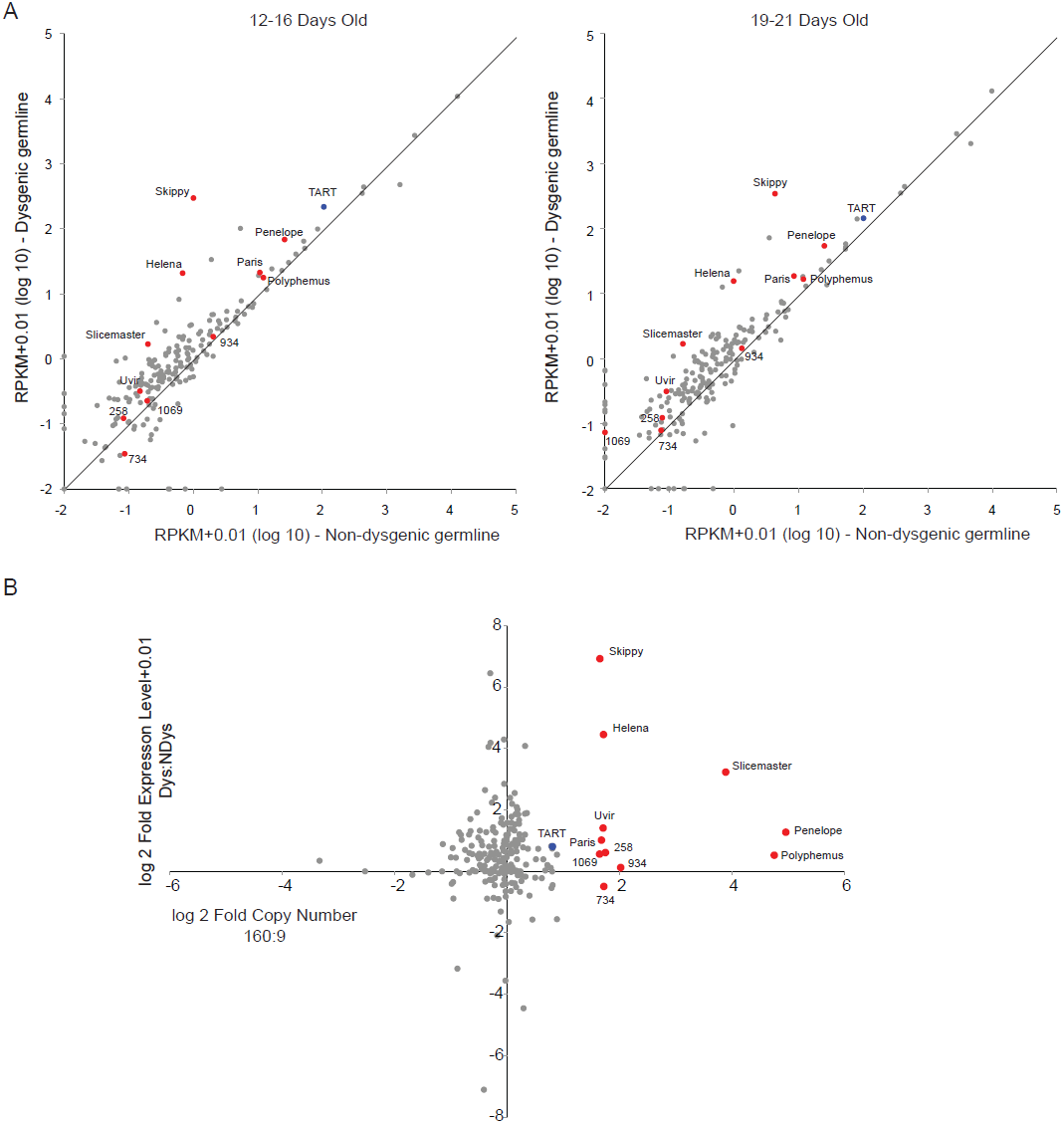
Increased TE expression in the dysgenic germline persists through adulthood. (A) RPKM + 0.01 for TEs (log 10), Dysgenic vs. Non-dysgenic germline. TEs that are in excess in 160 are more highly expressed, as well as many TEs that are not in excess. This state of increased global TE expression persists through adulthood. (B) Fold excess in expression (RPKM + 0.01,log 2, average between both ages) vs. fold excess in abundance in strain 160. Nearly all TEs that are in excess in 160 show increased expression in the dysgenic germline (11/12). But many TEs that are equivalent in abundance between strains are also increased in expression.

We next sought to determine whether the degree of expression in dysgenic progeny was largely determined by differences in bulk TE abundance between the two strains (Figure 2C). Ten of eleven elements that are more abundant in the inducer strain were expressed at higher levels in dysgenic progeny. However, many upregulated elements are similar in abundance between strains. In fact, many elements that are *more* abundant in strain 9 are upregulated in dysgenesis (Figure 2C). This includes the three elements that are not lowly expressed.

### Global Genic Mis-expression in a Dysgenic Cross

To determine how global TE derepression might modulate genic expression, we analyzed endogenous gene expression within the same 0-2 hour old embryos. There was a general increase in expression for lowly expressed genes (<5 RPKM) in the F1 progeny of the dysgenic cross. This is most easily demonstrated by observing relative expression level as a function of rank expression level genome wide (Figure 3). This coincided with a modest decrease in expression of the most highly expressed genes (Figure 3, right panel). Since RPKM is a proportional measure, a change in expression in highly expressed genes is expected to increase lowly expressed genes, and vice versa. Thus, it appears that the global distribution of gene expression is tilted in the dysgenic germline. To determine if this result could be explained by differences in the expression of genes primarily residing in heterochromatin, we excluded putative heterochromatic scaffolds smaller than 4 million base pairs and excluded genes residing closer than 1 million base pairs from either end of the remaining scaffolds. To further ensure this result was not driven by TEs that were computationally misidentified as genes, we eliminated all genes lacking orthology to *D. melanogaster*. Even after accounting for these effects, the trend is maintained (Supplemental 2), and is observed in technical replicates within biological samples as well as in flies of different ages.

**Figure 3.**
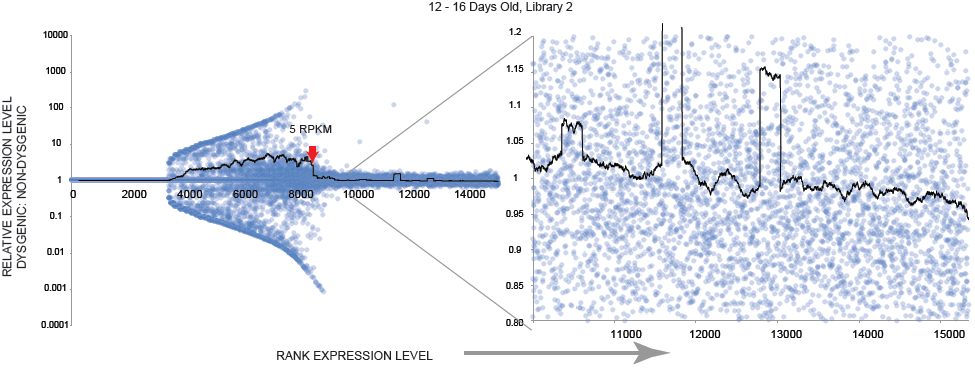
A shifted balance of gene expression in the dysgenic germline: lowly expressed genes are increased in expression in dysgenesis, highly expressed genes are decreased. Relative expression of endogenous genes (RPKM + 0.01, Dysgenic vs. Non-dysgenic) ranked from lowly to highly expressed genes for one mRNA-seq library (12-16 days old females, Library 2). A sliding window for the average ratio of sets of 250 genes is shown, showing an excess of lowly expressed genes that have higher relative expression in the dysgenic germline. Arrow indicates the 5 RPKM boundary of expression. Concordant with this, the same size sliding window (right panel) shows very highly expressed genes have reduced expression in the dysgenic germline.

Additionally, we performed differential gene expression analysis based on two metrics. We considered both fold-change in expression as well as p-values for differential expression by DESeq2[33] and edgeR[34,35]. Based on these metrics, we performed gene ontology enrichment analysis using GOrilla [36,37], which performs functional enrichment analysis based on ranked lists rather than cutoffs (Supplemental File 3). When sorted by fold expression level, we found that genes involved in positively regulating transcription are significantly enriched among genes more highly expressed in the dysgenic germline, consistent with our observed upregulation of many genes with dysgenesis. Among genes down regulated, genes involved in chitin synthesis were significantly enriched. However, clustering of chitin related genes within the genome causes a lack of independence among these genes. When sorted by p value, there is no enrichment of classes among genes upregulated in dysgenesis, perhaps arising from low statistical power for genes with low expression. For genes downregulated in dysgenesis, we again find that genes in the chitin synthesis pathway are down regulated. In addition, we find that many highly expressed ribosomal proteins as well as members of the SWI/SNF family of chromatin remodelers are enriched for down regulation in dysgenesis. In the former case, this is consistent with results showing a shift toward reduced expression among highly expressed genes.

### Germline transposable element piRNA in a dysgenic cross

To determine how maternal inheritance of piRNAs might explain persistent TE mis-expression in the dysgenic germline we sequenced piRNAs (defined as small RNAs, 23-30 nt, filtered against known non-piRNA classes) from 0-2 hour old embryos laid by strain 9 and strain 160 mothers. This allows a measure of maternally provisioned piRNA load. In addition, we sequenced piRNAs in embryos laid by F1 females from both reciprocal cross directions between the two strains to determine whether perturbed piRNA biogenesis during dysgenesis might maintain increased TE expression. For F1 germline piRNAs, we collected embryos from the same pool of mothers used for mRNA-seq, but at intermediate maternal age (15-16 days old). This allowed us to determine whether the persistent mis-expression of TEs in the F1 germline of the dysgenic cross could be explained by a persisting defect in piRNA biogenesis.

Figure 4A shows that there is a large number of TEs, including many of those with greater copy number in Strain 160, with higher levels of 160 maternally provisioned piRNA. Strikingly, despite the asymmetry in maternal provisioning, piRNA levels largely equilibrated in reciprocal F1 germlines (Figure 4B). A significant exception to this is the *Helena* element.

**Figure 4.**
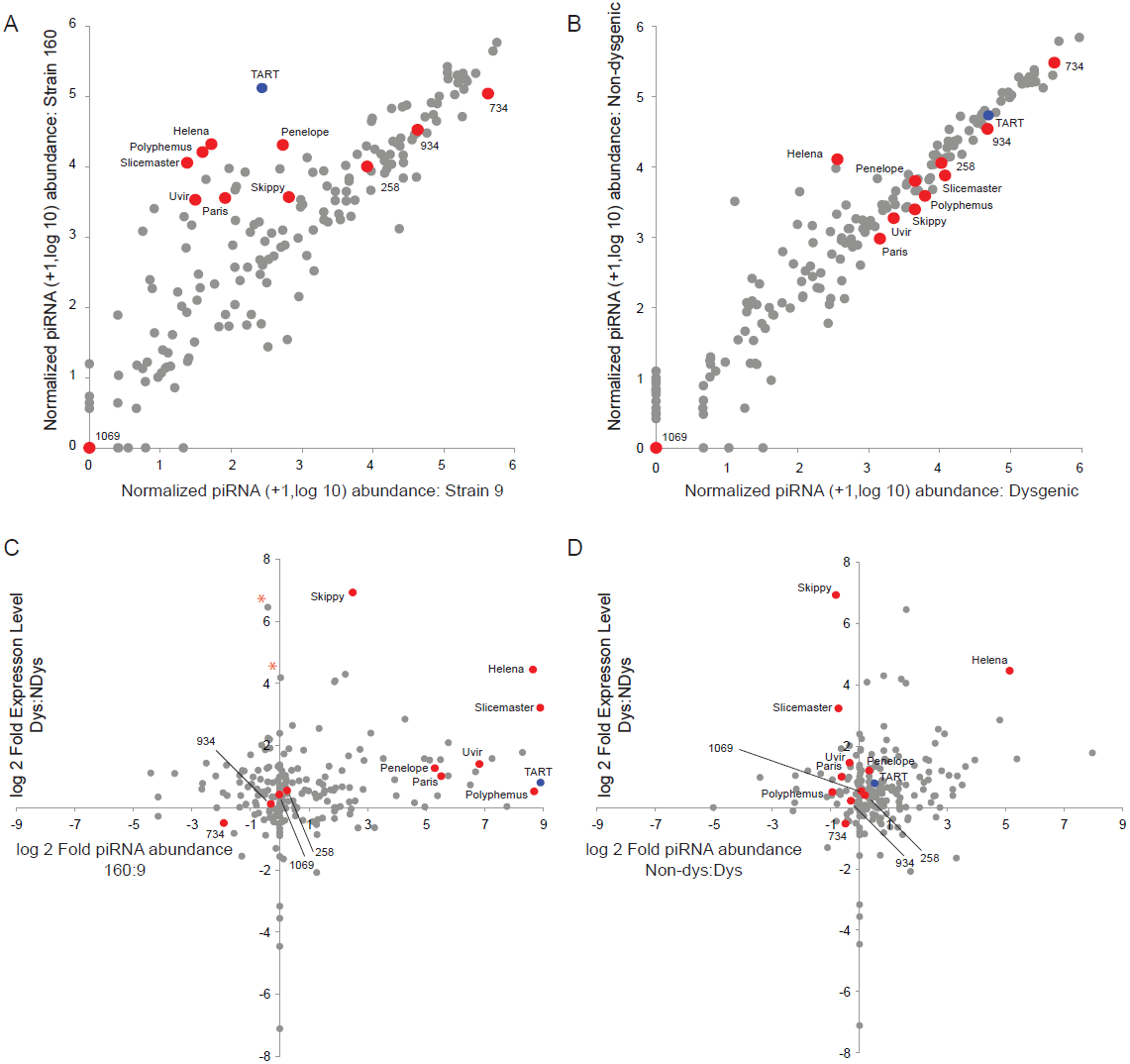
Despite persisting TE overexpression, piRNA abundances are largely equilibrated in the dysgenic germline. (A). Normalized (per 10 million mappers) piRNA abundance + 1 (log 10) in the strain 9 germline vs. the strain 160 germline. A large number of TEs show increased piRNA expression in the strain 160 germline, especially TART and others enriched in abundance in strain 160. (B) Normalized (per 10 million mappers) piRNA abundance + 1 (log 10) in the dysgenic germline vs. the non-dysgenic germline. piRNAs abundances for many TEs with greater excess in strain 160 become equilibrated in the dysgenic germline. A significant exception to this is the *Helena* element. (C) TE piRNA excess in strain 160 vs. relative expression level in dysgenesis. TE piRNA asymmetry between 160 and 9 is not the sole determinant of increased expression in dysgenesis. Some TEs, such as those denoted with (*), are increased in expression in dysgenesis, despite similar piRNA abundances in 9 and 160. (D) TE piRNA excess in the non-dysgenic germline vs. relative expression level in dysgenesis. Many TEs with increased expression in dysgenesis are equilibrated with respect to piRNA abundance, though this is not the rule. *Helena* stands out as a strong exception.

Figure 4C demonstrates the degree to which asymmetry in maternal provisioning predicts TE expression in reciprocal progeny in the next generation. Many of the elements that are more abundant in strain 160 have greater piRNA abundance in the 160 female germline and nearly all of these show greater expression in the germline of the dysgenic cross. However, at least two elements that are overexpressed in the dysgenic germline (indicated with a *) show equal or higher piRNA abundance in the 9 germline. The second most differentially expressed element in the dysgenic germline shows little difference in piRNA abundance between strains. Therefore, maternal provisioning of piRNA is an imperfect global predictor of TE expression in hybrid dysgenesis.

To determine whether this chronic level of TE mis-expression might be explained by persistent asymmetry in germline F1 piRNA pools, we investigated the degree to which asymmetry in F1 piRNA levels was predictive of expression in reciprocal progeny. Overall, there is a poor relationship between relative piRNA abundance for a given TE and its relative expression difference in dysgenic compared to non-dysgenic germlines. Furthermore, there is a contrasting pattern within the class of TEs that are higher in copy number in 160 and also more highly expressed in the dysgenic germline. In particular, two elements that become overexpressed during dysgenesis (*Skippy* and *Slicemaster*) in fact have higher piRNA abundances in the dysgenic germline. Thus, it is clear that piRNA biogenesis is maintained, but is not sufficient to maintain TEs at a low level of expression. *Helena* is the only element more abundant in strain 160 (at a 3X level) whose increased expression level in dysgenesis is explained by reduced levels of F1 piRNA.

Raw abundance measures of piRNAs ignore critical aspects of their biogenesis and two recent studies have demonstrated that globally reduced signatures of robust piRNA biogenesis likely contribute to the mobilization of diverse TEs [21,38]. Overall, we see no evidence that piRNA biogenesis is skewed based on the size distribution of small RNA reads (Figure 5A). For each TE we estimated the percent ping-pong[20] as well as the density of ping-pong pairs in dysgenic and non-dysgenic germline. When comparing metrics directly, there is little evidence that piRNA biogenesis is grossly perturbed in the dysgenic cross though a more sensitive comparison using normalized Z-scores indicates a modest shift in piRNA abundance beyond what would be expected based on patterns of maternal provisioning (Figure 5B). This is observed in the z-score heat maps for abundance and ping-pong pair density, which show an excess of negative z-scores for dysgenesis (p<0.0001 for both differences, Wilcoxon Signed-Rank Test). Importantly, these are both normalized, proportional measures of abundance that are likely influenced by abundance of genic piRNAs in the same library (see below).

**Figure 5.**
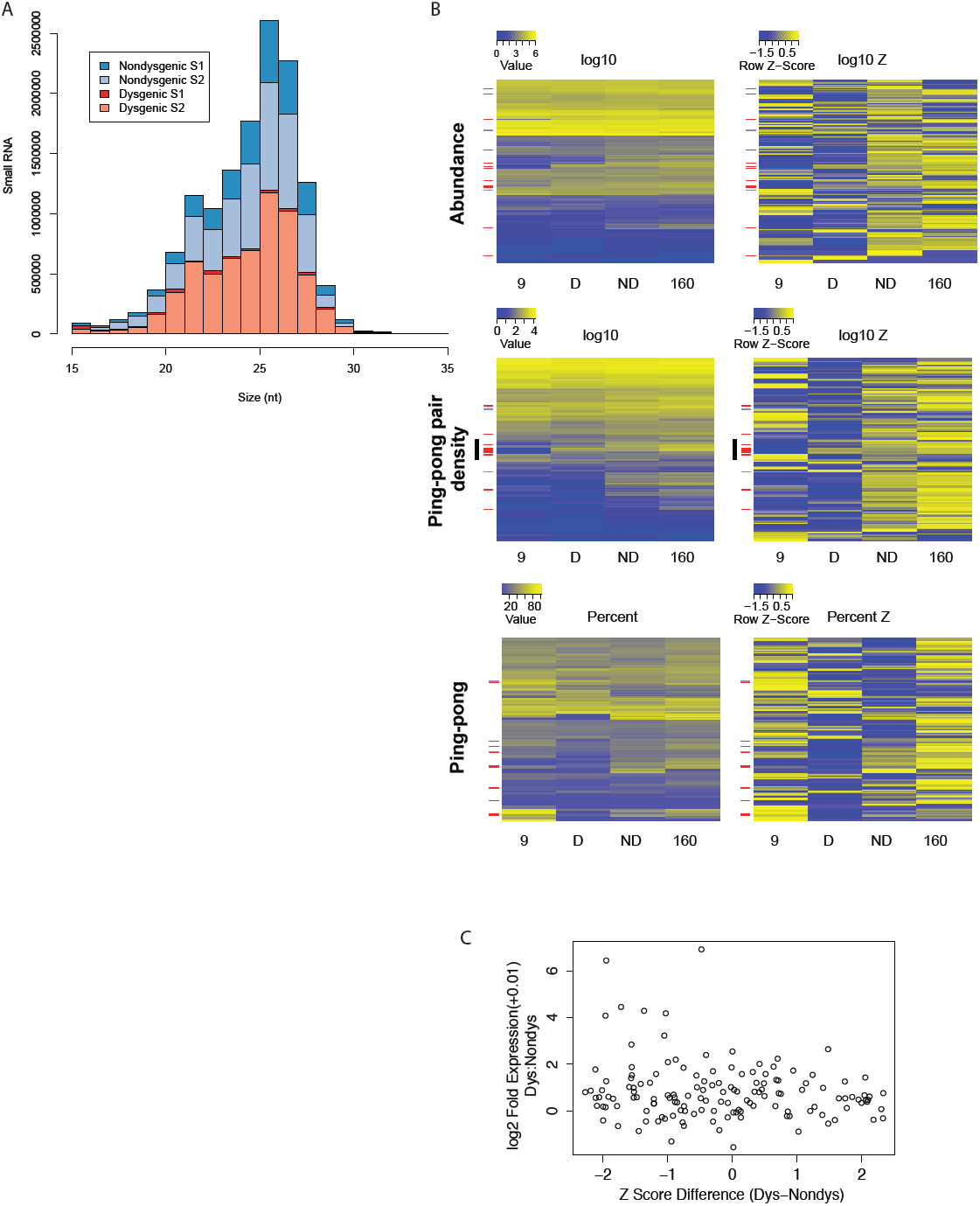
Signatures of piRNA biogenesis in the dysgenic germline show only modest defects. (A) Size distributions of small RNAs are similar between dysgenic and non-dysgenic germlines. Distribution of all small RNAs (not normalized) from four germline libraries (2 dysgenic, 2 non-dysgenic) filtered for tRNA, rRNA and snoRNA. (B) piRNA biogenesis signature heatmaps. TEs upregulated in the dysgenic germline (a difference of 5 RPKM or higher) are indicated with red bars. TEs upregulated in the non-dysgenic germline (a difference of 5 RPKM or higher) are indicated with purple. On the left are heatmaps for raw measures of abundance, the density of ping-pong pairs and percent ping-pong. On the right are heatmaps for the same metrics, but by row z-score. For raw measures, there are no noticeable effects of dysgenesis on piRNA biogenesis. Row z-scores in dysgenesis do show lower values for abundance measures (abundance and ping-pong pair density), but not percent ping-pong (see text).(C) Fold excess in expression in dysgenesis vs. the difference in percent ping-pong between dysgenic and non-dysgenic. Of the top eight that are most differently expresses in dysgenesis, all have lower ping-pong z-scores in dysgenesis.

In contrast to these abundance measures, there is no evidence for global ping-pong biogenesis disruption as measured by percent ping-pong. Note, for example, that many row-z scores for percent ping-pong show weaker z-scores for non-dysgenic compared to dysgenic (compare upper and lower portions of the percent ping-pong z-score heatmap). A Wilcoxon Signed-Rank test also found no significant difference (p=0.22) in percent ping-pong between dysgenesis and non-dysgenesis. Thus, in contrast to previous studies, we find no evidence for a global disruption of piRNA biogenesis.

Despite this, one can see that elements that are overexpressed in dysgenesis (red bars) are in excess among elements with a reduced ping-pong signature (bottom portion of percent ping-pong z-score heatmap). Furthermore, there is a slightly significant correlation between difference in percent ping-pong z-score and fold expression (p=0.048, Figure 5C). This trend is driven by the top eight elements that show higher expression in dysgenesis and all have lower z-scores in dysgenesis. Overall, this does not support a model of global disruption in piRNA biogenesis maintained in adult flies. Rather, the persistence of higher expression for some TEs is driven by idiosyncratic defects in piRNA biogenesis that occur on a TE-by-TE basis.

### Genic piRNAs in hybrid dysgenesis

Recent studies have demonstrated that in addition to TEs, genes may also be the target of piRNA silencing. Genic targeting by piRNAs can arise from neighboring TE insertions that drag flanking sequences into piRNA biogenesis and gene silencing in *cis* [39,40,41,42,43]. Previous important work in the *D. virilis* system of dysgenesis identified a piRNA cluster overlapping the *center divider* (*cdi)* gene (Dvir\GJ14359) [44]. We thus sought to determine how the global landscape of genic piRNAs was influenced by the activation of diverse transposable elements. Global gene expression might be modulated if genic piRNA silencing were either attenuated or enhanced during dysgenesis. To avoid gene models that might designate TEs to be genes, we only examined the CDS regions of *D. melanogaster* orthologs.

In the dysgenic germline the number of genes being processed into piRNAs was significantly higher than in non-dysgenesis. Figure 6A indicates all CDS’s that have at least 50 piRNAs per 10 million mapped, row normalized by z-score. A very large number in dysgenesis show a unique signature of being targeted by piRNA and these *de novo* targets largely become a source of sense only piRNAs (Figure 6B). The genes that showed a significant increase in piRNA were also more highly expressed above the genome wide background (Figure 6C, Wilcoxon rank test, p<0.0001) and tend to become more lowly expressed in the dysgenic germline compared to the non-dysgenic germline (Figure 6D, Wilcoxon rank test, p<0.0002). GO enrichment analysis indicated that genes that specifically become sense piRNA biogenesis targets are enriched for highly expressed ribosomal protein RNAs (FDR p-value: 2E-08, Supplemental 3). This is concordant with mRNA-seq results from independent RNA collections that show that ribosomal protein genes are also enriched among down regulated genes in dysgenesis. The fact that these piRNAs are predominantly sense suggests that the observed modest levels of reduction in gene expression for highly expressed ribosomal genes may be explained by being processed into piRNA biogenesis in an off-target manner without a corresponding anti-sense pool that amplifies the off-target silencing. One possibility is that the mechanism underlying perturbed piRNA biogenesis for only a subset of TEs also drives genic off-targetting of highly expressed genes. This may occur through cytoplasmic, non-specific slicing of mRNAs that are highly abundant.

**Figure 6.**
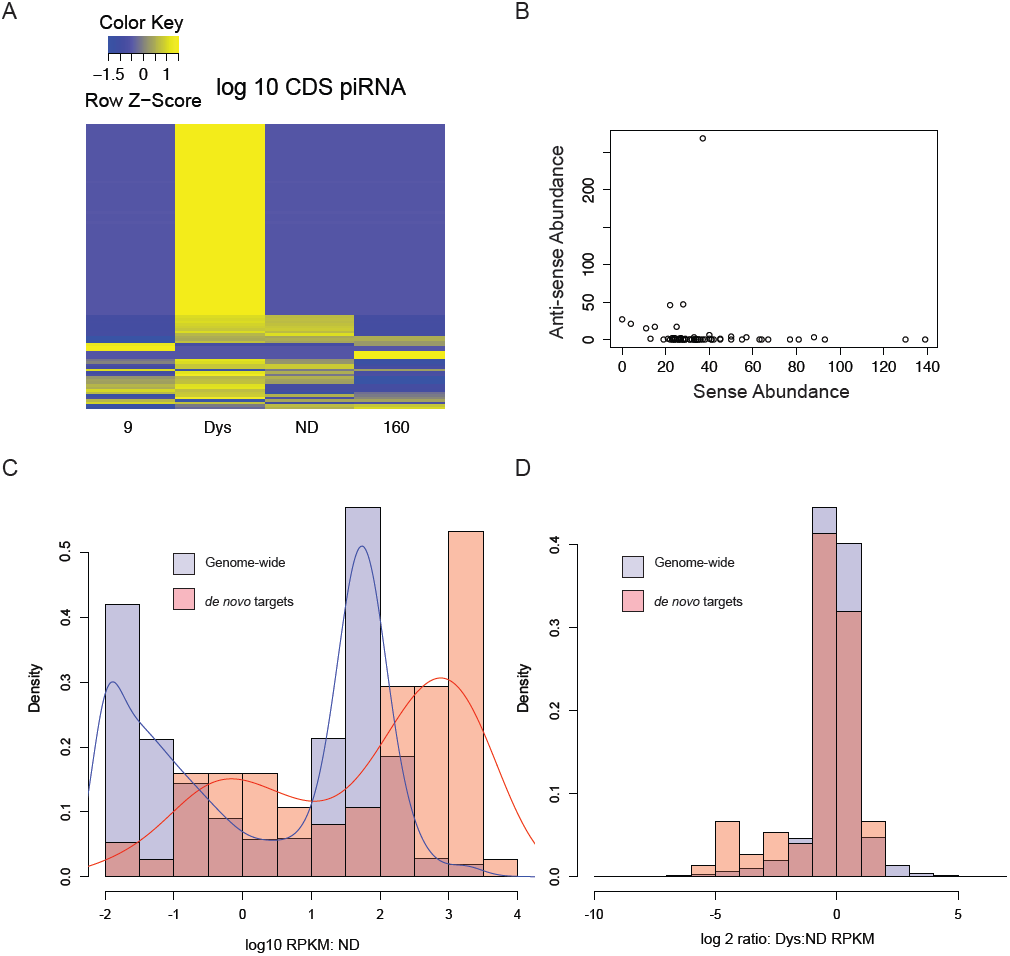
Highly expressed genes are a *de novo* source of sense-piRNAs in the dysgenic germline. (A) Heat map of *D. melanogaster* ortholog CDS regions that are piRNA targets (above a threshold of 50 piRNAs per CDS per 10 million mapped for at least one CDS). A large class of CDS are a *de novo* source of piRNAs in dysgenesis. (B) Sense vs. Anti-sense abundance for piRNAs in the *de novo* class. piRNAs are highly biased toward the sense strand. (C) Distribution of expression levels (log 10 RPKM) for *de novo* targets vs. all genes in the genome. *de novo* targets are more highly expressed. (D) As a source of sense piRNAs in dysgenesis, the expression of these highly expressed *de novo* targets is reduced. Shown is the distribution of expression ratios (dysgenic:non-dysgenic) for all genes and *de novo* targets.

### Genetic Analysis of the Cause of Hybrid Dysgenesis

We have shown that hybrid dysgenesis in *D. virilis* is associated with a persisting global shift in both TE and gene expression. We next sought to identify the genomic regions of strain 160 that most protect against hybrid dysgenesis in a cross with 160 males. Excess TART abundance, as well as previous reports of unique genic cluster behavior at chromosome ends [44], indicate a possible role of telomeres. Since maternal repression of dysgenesis is a dominant trait, we diluted fragments of the 160 genome onto a strain 9 background though backcrossing hybrid females to strain 9 males. Two generations of backcrossing through females allowed multiple generations of recombination in the absence of dysgenesis. We then tested F3 females for protective ability by crossing to inducer males, followed by genotyping at markers distributed across the genome. In particular, each F3 chromosome was genotyped for a SNP marker near the telomere, near the centromere and also along the euchromatin of the chromosomal arm.

With this backcross scheme, we generated a panel of females with significant variation in their ability to protect against induction when crossed to 160 inducer males (Figure 7A). QTL analysis identified three genomic regions from strain 160 that significantly protected against F1 sterility (Figure 7B) The region most significantly protective corresponded to the pericentric region of chromosome 5. The pericentric region of the X chromosome was also significantly protective, followed by a euchromatic region in the proximal arm region of chromosome 4. A previous study indicated that the most protective chromosome was also chromosome 5, followed by the X and then chromosome 4 [32]. Interestingly, the most protective region of the *D. virilis* genome corresponds to the pericentric region of the same Muller element that carries cluster 42AB in *D. melanogaster*. This provides strong genetic evidence for a role of pericentric, cluster-derived piRNAs in the protection against dysgenesis in *D. virilis*. By contrast, no telomeric region provides significant protection. Thus, amplified TART elements seem to be a passenger of TE destabilization in strain 160 rather than a driver.

**Figure 7.**
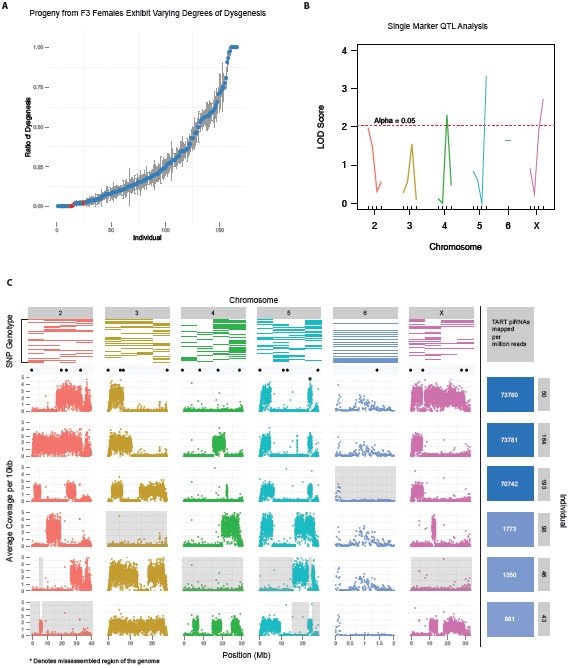
No single genomic region is necessary for maternal protection against dysgenesis, but pericentric regions are the strongest. (A) Scatterplot showing proportion of dysgenic testes (*y* axis) observed in the progeny of each F3 female individual (*x* axis). Red dots indicate F3 females that were selected for whole genome sequencing. (B) Single marker QTL analysis identified 3 putative QTLs: one flanking the centromeres of the 5^th^ and X chromosomes and one of the tested euchromatic regions of the 4^th^ chromosome. (C) Top row: Results from the genotyping assay. Colored rectangles represent the presence of strain 160 SNPs in individuals, ranked from top to bottom (most protective individuals on top). Scatterplots: sequencing results. Each dot represents the average number of base pairs that uniquely mapped to every 10 kb of the 160 genome. Valleys indicate regions of 9 homozygosity. Black dots above scatterplots show the location of each SNP used for our genotyping assay. Grey background demonstrates that no region of the genome from 160 is necessary or sufficient to protect against dysgenesis. Right-most column: Number of piRNAs mapped to TART sequences, per million reads, for each F3 female individual. Color intensity is representative of TART piRNA abundance.

In addition to standard QTL analysis, to determine precise regions of the 160 genome critical for protection, we performed whole genome sequencing of the top 6 most protective females for which DNA was available (Figure 7C). A synthetic diploid genome sequence was generated from both strain 160 and strain 9 haploid consensus sequences and reads that strictly mapped uniquely to the strain 160 haploid genome were identified. This allowed us to identify regions of heterozygosity for strain 160 over a strain 9 background for the six most protective mothers. This provided maximal resolution for a test to rule out whether any single genomic region is critical for maternal protection. From this analysis, we found that among the most protective mothers there is no single genomic region of strain 160 always present in each most protective mother. Across all six mothers, we identified at least one mother homozygous for strain 9 at each position of the genome (excluding unassembled regions). Thus, no single genomic region appears critical for protection against dysgenesis.

To determine whether piRNA from any particular TE enriched in 160 was dispensable for protection, we sequenced small RNAs from the ovaries of the six most protective mothers. We first reasoned that the only TEs that are candidate inducers of dysgenesis are those for which piRNA abundance is greater in strain 160 than in strain 9. Among the 221 repeats within the library, there are 141 that meet this qualification. These are candidates for elements contributing to the dysgenic syndrome. We further reasoned that if any female with full repressive ability had piRNA abundances for a TE that was similar to strain 9, we could also rule out that TE as a driver of dysgenesis. Of the remaining 141 candidates, 88 have at least one protective female that has normalized piRNA abundance less than or equal to strain 9. Thus, we are left with 53 candidate contributing repeats. Using these criteria, we were unable to eliminate any of the elements 3-fold enriched from strain 160. However, the data further demonstrate a minimal role for the TART element piRNAs as mediators of maternal protection. This is because the vast majority of TART piRNAs in the protective mothers are derived from the tip of the X chromosome from 160 (Figure 7C) and three of the six females that are most protective against dysgenesis lack the X 160 telomere allele. Notably, the telomeric region of the X has special silencing properties in *D. melanogaster* that may explain the excess of TART piRNAs derived from this one genomic region [45,46,47,48,49], but in this system this region plays no role in protection against dysgenesis.

### Genic piRNA cluster behavior across generations

A previous study by Rozhkov *et al*. demonstrated that strain 160 possesses a number of piRNA clusters that are absent in strain 9 [50]. This may be related to the fact that strain 160 also carries many TEs in greater copy number compared to strain 9. Additionally, it was noted that many of these clusters were telomeric which is consistent with results here that show a signature of increased TART activation in strain 160. The 160 and 9 strains in our lab have been separated from these strains for more than 15 years, so this allows a comparison of cluster properties among strains of flies that have been long separated. The most well-characterized cluster from Rozhkov *et al*. was a telomeric cluster at the tip of chromosome 2 encompassing the gene *center divider (cdi)*. This dual strand piRNA cluster was only in strain 160 and absent in strain 9 and this pattern was also observed in our divergent laboratory stocks (Figure 8). Interestingly, a second cluster identified by Rozhkov *et al*, near the telomere of the 6th chromosome in strain 160, but absent in strain 9, showed an opposite pattern in our strains. This cluster in our strain 160 had 318 unique mappers per million mapped whereas our strain 9 had 3,836 unique mappers per million mapped. Rather than being independently gained twice, the most parsimonious explanation for this was that this cluster was originally in both lines, but independently lost in our strain 160 and their strain 9. Thus, some piRNA clusters may be prone to losing their activity over time.

**Figure 8.**
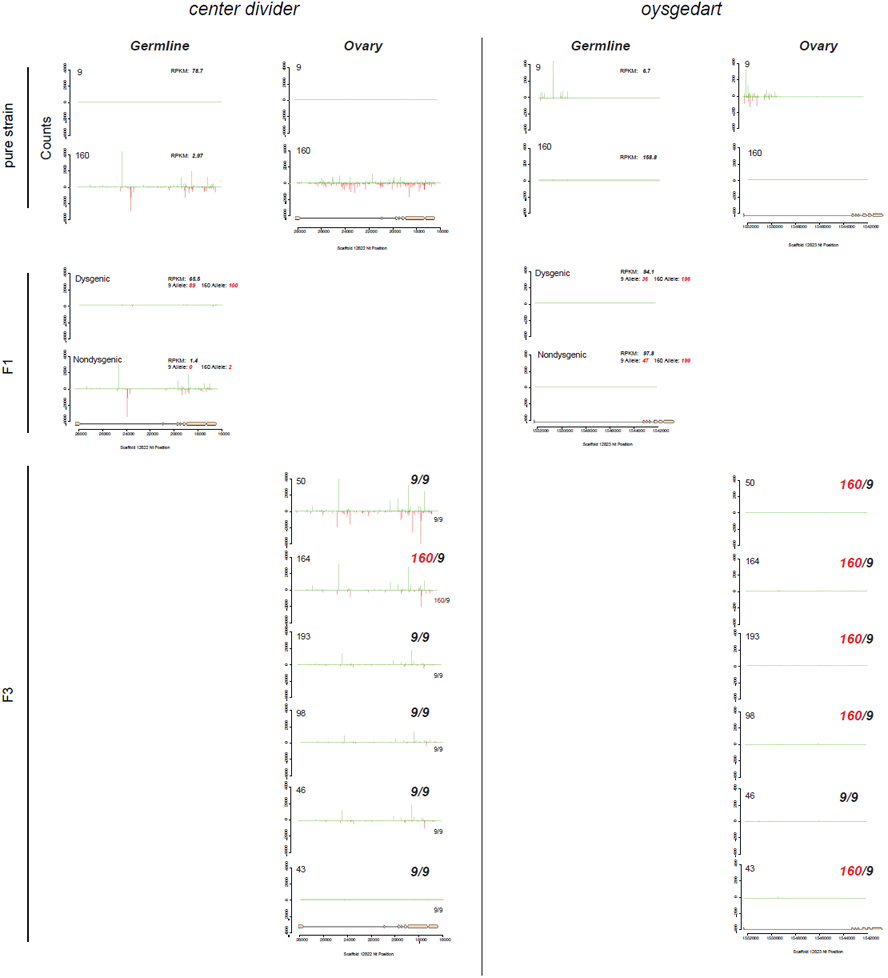
Germline and ovary genic cluster behavior across generations for *D. virilis* orthologs of *center divider* and *oysgedart* from *D. melanogaster*. piRNA mapping densities are indicated. mRNA-seq RPKM for germline (0-2 H embryo) is also indicated. Allelism was determined by counting mRNA-seq reads based on SNPs that distinguish strain 9 and 160. Strain 160 cluster identity is maintained for *cdi* in non-dysgenic progeny in which strain 160 is the mother. This is correlated with silencing of both alleles in the non-dysgenic germline. In contrast, the cluster is not maintained in the dysgenic germline and both alleles are expressed. Germline cluster identity for *oysgedart* (which in the germline is predominantly sense) is lost in progeny. In this case, expression is even between reciprocal progeny, but germline expression is lower off the 9 allele in both directions of the cross For cluster behavior in F3 backcrosses, heterozygosity or homozygosity of the respective allele is indicated. Notice how cluster identity is maintained for *cdi* to varying degrees in individuals homozygous for the 9 allele. Cluster identity may be strongest in lines carrying the high TART piRNA abundance allele at the tip of the X chromosome, suggesting cluster activation in *trans*. In contrast, cluster activity is absent in all progeny for *oysgedart*.

Additionally, we identified the *oysgedart* gene (Dvir\GJ17620) in strain 9 that displayed a unique mode of genic piRNA targeting. A *Ulysses* element insertion upstream of this gene in strain 9 is specifically associated with genic silencing and shunting *oysgedart* into piRNA biogenesis (Figure 8). piRNAs derived from *oysgedart* are dual-strand in strain 9 ovaries (which includes somatic and germline tissue) but biased towards the sense strand in the germline.

To determine how different modes of genic targeting by piRNA influences gene expression across generations, we examined patterns of gene expression in the germline in strain 9 and 160 parents (from 0-2 hour embryos), as well as reciprocal offspring. For reciprocal hybrid offspring, we also examined patterns of somatic gene expression in carcasses with ovaries removed to determine how maternally provisioned piRNA might influence somatic expression in the next generation. Finally, we examined cluster behavior in the F3 backcrosses (using ovary piRNA data) to determine how genic clusters are maintained when the original cluster allele is segregated away from a non-cluster allele.

We found that two modes of genic targeting by piRNAs result in contrasting behavior across generations. The *cdi* dual strand cluster is maintained as a source of germline piRNAs when it is inherited from the mother (Figure 8). However, when inherited paternally, the cluster is inactive in the germline. In addition, *cdi* expression is highly reduced in the germline of strain 160 and the germline silencing of both alleles of *cdi* is maintained when transmitted maternally and made heterozygous in combination with the wild-type strain 9 allele. In contrast, when the *cdi* cluster allele is transmitted paternally, piRNA cluster activity is reduced and germline expression levels are maintained near the level of strain 9, with similar contributions from each allele. This asymmetry in gene expression of *cdi* (off in non-dysgenic, on in dysgenic), however, is not observed in the soma. RNA collected from carcasses of F1 females, with ovaries removed, of dysgenic and non-dysgenic crosses (3 individual carcasses performed separately from each direction), showed similar levels of expression from both directions of the cross (Dysgenic Carcass: 17.1 RPKM [9 Allele: 7 counts;160 Allele: 9 counts] vs. Non-dysgenic Carcass: 23.4 RPKM [9 Allele: 5 counts; 160 Allele: 12 counts]) Therefore, silencing of germline *cdi* occurs in *trans* when the cluster is inherited maternally, but this has no effect on the somatic expression.

The *oysgedart* allele in strain 9, being silenced by off-target processing into primarily sense piRNA in the germline, provides a significant contrast. In neither direction of the cross is this form of sense piRNA biogenesis maintained in progeny. Instead, the wild-type allele from strain 160 seems to function in *trans* to limit this mode of silencing. Thus, in both directions of the cross, the expression of *oysgedart* is maintained, but only at about 60% wildtype level, since the *Ulysses* insertion allele is more lowly expressed. While sense piRNA biogenesis is turned off in both reciprocal directions, expression of the 9 allele is always reduced in *cis*. Interestingly, in neither direction of the cross does there appear a maternal effect on somatic expression for *oysgedart*. Even though expression of the 9 allele is reduced in the germline, it remains on in the soma of both dysgenic (RPKM: 62.0, [9 Allele: 21 counts; 160 Allele: 21 counts]) and non-dysgenic (RPKM: 80.9, [9 Allele: 29 counts; 160 Allele: 25 counts]) F1 females. This also demonstrates that the allelic *cis* effects on local silencing by the Ulysses insertion are germline, not soma, specific (Fisher’s exact test for difference in allele effects between soma and germline. Dysgenic: p<0.0001, Nondysgenic: p<0.0001)

Using the genome and small RNA data from the six females used for genetic analysis, we were also able to observe the behavior of the *cdi* cluster across multiple generations in cases where the original *cdi* dual strain cluster allele from 160 had been segregated away and only a homozygous strain 9 allele was present. From this analysis, the *cdi* cluster demonstrates the ability to modestly paramutate piRNA biogenesis onto the strain 9 allele. To varying degrees, cluster behavior was maintained in most flies homozygous for the strain 9 allele of *cdi*. In one case, cluster behavior was lost. Because the cross scheme was maintained over two generations, we are unable to determine the generation in which the cluster activity is lost. However, it is clear that *cdi* can paramutate in one generation but that this paramutation is not robust across multiple generations. This was never observed for *oysgedart*.

## Discussion

Females that are the product of reciprocal crosses are genetically identical but differ from each other epigenetically due to parental genotypic effects. Hybrid dysgenesis represents an extreme form of phenotypic variation among genetically identical individuals. Here we show that differences between genetically identical dysgenic and non-dysgenic individuals is manifested in multiple ways. When genomes from two strains of *D. virilis* are brought together, TEs that are more abundant in one genome become more highly expressed in the germline of the next generation. Coincident with this, there is a persisting increase in TE expression for several TEs that are evenly distributed between strains. This can be explained by defects in piRNA biogenesis that occur on a TE-by-TE basis. In addition, there is a global shift in the distribution of endogenous gene expression. Our results suggest that changes to endogenous gene expression are, in part, a consequence of *de novo* genic piRNA production in dysgenic compared to non-dysgenic progeny. Highly expressed genes become off-targets for piRNA biogenesis and their expression is reduced. The mechanism for this is unclear, but may be driven by the same mechanism that leads to idiosyncratic defects in piRNA biogenesis for some TEs.

We identify a large number of differences in TE expression, genic expression, TE-derived and genic-derived piRNA biogenesis between dysgenic and non-dysgenic progeny. Therefore, it is difficult to distinguish between causal factors and downstream effects. However, our genetic analyses clearly demonstrate that both induction of and protection against dysgenesis is distributed across the genome. Since multiple TE families are in excess copy number in the inducer strain, the weight of evidence favors a complex mode of hybrid dysgenesis driven jointly by multiple elements. This is supported by the fact that the regions of the genome that are most protective against dysgenesis are located in the pericentric regions, which are known to be critical sources of piRNAs [51].

In principle, there may be multiple mechanisms underlying the effects of dysgenesis on TE derepression and increased genic targeting. This is because other maternal effects are evident in this system. In particular, germline expression of *cdi* differs between dysgenic and non-dysgenic progeny. Differences in levels of expression of this gene, a tyrosine-protein kinase, could mediate cascading effects in different ways. Alleles of *cdi* share properties with imprinted genes since expression depends on which parent the allele is inherited from. However, *cdi* differs from canonical imprinting in that a silenced *cdi* allele is capable of silencing the other allele in *trans* when inherited maternally, similar to paramutations observed in maize, mice and recently in *Drosophila melanogaster* [52,53,54,55]. Imprinted genes can have a significant downstream effect on patterns of gene expression [56]. Even in *Drosophila*, where there is no evidence for DNA methylation, there are significant, albeit poorly understood, parent-of-origin allelic effects on global gene expression [57,58].

Two recent studies have identified globally defective piRNA biogenesis as a cause for global TE mis-regulation in hybrids between genetically divergent strains (*e.g.,* the P-element system in *D. melanogaster* and hybrids between *D. melanogaster* and *D. simulans*) [21,38]. In the *D. virilis* system studies here, differences in maternal piRNA provisioning likely explain the activation of multiple TEs in dysgenic progeny germline. However, we only find modest evidence that defects in piRNA biogenesis contribute to global TE mis-expression with dysgenesis. Even while TE mis-expression persists, piRNA abundance is largely equilibrated between reciprocal progeny, with the exception of the *Helena* element. Only modest defects are observed in piRNA biogenesis signatures and these perturbances only pertain to some TEs.

Related to this point, there are several critical differences between this syndrome of dysgenesis and the *D. melanogaster* P element system that has been most thoroughly characterized. First, the induction of gonadal atrophy is nearly 100% at the sensitive temperature in the P element system. In the *D. virilis* system, complete sterility is observed in approximately 50% of F1s, and the sterility effect has ameliorated since it was first observed[10]. Second, near complete P element mediated sterility is proposed to be gradually rescued by new TE insertions into piRNA clusters as flies age[21]. These new insertions establish a new pool of protective piRNA that in turn lead to restored TE repression. A significant difference between the P element system and the *D. virilis* system is with respect to the ability of F1 females of the dysgenic cross to again protect in a second generation cross. In *D. virilis*, near complete protection against dysgenesis is present in heterozygous females that have inherited all 160 chromosomes paternally [32]. Thus, the ability to transmit protection against dysgenesis is restored in one generation and this is born out by the fact that piRNA pools are largely equilibrated in reciprocal progeny. Functional restoration of maternal protection in the *D. virilis* system argues strongly against a persistent global defect in piRNA biogenesis in the adult F1 germline. Importantly, many of the elements that are in excess in 160 do in fact exist as older variants in strain 9. piRNAs targeting these older TE variants are present in strain 9, albeit at a lower level than in strain 160. Previous studies have demonstrated that older divergent piRNA pools play an important role in re-establishing control of re-proliferating TEs [18]. By contrast, multiple generations of female P chromosome transmission are required to re-establish the P cytotype after one generation of paternal transmission [59].

It has recently been proposed that parent-of-origin effects made evident from reciprocal F1 crosses may make an important contribution to heritability [56]. These effects are mediated through a regulatory network via imprinted genes. Here, using a system of hybrid dysgenesis in *D. virilis*, we show that parent-of-origin effects can influence germline integrity but also influence the expression level of many genes through numerous indirect mechanisms. It will be interesting to determine whether early events in germline establishment mediated by parent-of-origin effects can lead to persistent differences in gene expression among individuals in natural populations.

## Methods

### Custom *D. virilis* TE library

Few annotated TEs are available for *D. virilis*. Therefore, we combined available annotated TE sequences with two computationally predicted libraries (generated with PILER [60] and REaS [61,62]

ftp://ftp.flybase.net/genomes/aaa/transposable_elements/) to generate a manually curated library. Several annotations (*Uvir, Helena, TART, Telemac*) were improved by manual curation. Portions of the PILER library were also manually curated. Additional sequence from the *Helena* element was obtained by interrogating a *de novo* assembly of the strain 160 genome. Redundancy was removed from this combined library first by removing repeats with significant blastn hits between and within the PILER and annotated library, with priority to annotated and longer sequences. With this filtered set, further redundancy was removed by blasting this library with and between the REaS library.

### Genome Sequencing and TE measurement from strain 9 and 160

Wandering 3rd larvae were collected from strain 9 and 160, rinsed with 50% bleach and DNA was extracted. 100 bp paired end sequencing was performed on an Illumina GAII with 400 to 500 bp fragments. TE abundance estimation was performed with single ends that were trimmed by Sickle (https://github.com/najoshi/sickle), mapped to the TE library with BWA-MEM[29] and normalized by read numbers mapping the reference. Homogeneity within mapped reads was measured using piledriver (github.com/arq5x/piledriver) and averaging the frequency of the major allele across all nucleotides within the mapping for each TE.

### Estimation of zygotic effects of paternally inherited chromosomes

From the genome sequences of strain 9 and 160, restriction fragment length polymorphisms were identified that distinguish between the strains at two positions for each chromosome. F1 males were generated from a non-dysgenic cross and these males were crossed to strain 9. 96 F2 progeny were collected, (48 from each class, dysgenic or non-dysgenic) and genotyped for chromosomes inherited paternally with RFLPs. Log-odds ratios for the probability for being dysgenic with a given chromosome were estimated using a generalized linear model for logistic regression (binomial family with a logit link) in R. Some failed genotypes resulted in N=92.

### mRNA seq: Dysgenic and Nondysgenic germline, Strains 9 and 160 germline, dysgenic and non-dysgenic soma

RNA for sequencing was collected from embryos laid by F1 mothers from the dysgenic and non-dysgenic directions of the cross. Ovaries were not selected because dysgenic ovaries are often atrophied. Therefore, the germline tissue represented in this experiment is derived from mothers that have escaped germline ablation. Paternal effects on embryos that might occur when dysgenic females are mated with sterile dysgenic brothers were minimized by equally mixing males from reciprocal directions of the cross and allocating them in mating cages between reciprocal F1 females. This also ensured improved egg laying from dysgenic females. Females were maintained as continuously laying with a constant supply of yeast and grape plates and eggs were collected after 0-2 hour durations and flash frozen in liquid nitrogen. RNA was pooled from collections of 12 to 16 day old mothers and 19 to 21 day old mothers. From each biological age replicate, two RNA-seq libraries were generated for single-end, 50 bp sequencing, for a total of four libraries per condition (dysgenic or non-dysgenic). Reads were quality trimmed at the 3’ end (up to 16 bp) and reads with 2 bp of quality less than 20 were excluded using the Galaxy server. TE RPKM estimates were obtained by directly mapping with BWA to the annotated TE library and normalizing by mapping to the reference genome. mRNA RPKM estimates were obtained using the RNA-seq tool in CLC. Fold analysis was performed by calculating RPKM(+0.01) ratios. Ranking by p value was performed based on p values estimated with the DEseq2 and edgeR packages from raw counts. p value estimates are not true estimates since among the four libraries used for DEseq2 and edgeR analysis, there are only two biological replicates. However, p values were only used to estimate ranks for ontology enrichment.

Additional germline mRNA seq was performed using the same protocol for pure strain 9 and strain 160 (RPKMs averaged for cluster analysis across 2 libraries each for 7-15 day old females and 15-25 day old females). Allele counts were determined by direct counting within the RNA-seq mappings (summed across all library mappings) for a SNP known to distinguish the two strains within the transcripts for *cdi* and *oysgedart*. Somatic mRNA seq analysis was likewise performed, but from using RNA from dysgenic and non-dysgenic females with abdomens fully removed (3 libraries per condition, each library from asingle female, RPKMs averaged across libraries and allele counts estimated as before).

### Small RNA sequencing

All small RNA was size selected from 15% acrylamide, cut between an 18 bp oligo and the 30nt rRNA. Small RNA sequencing for dysgenic and non-dysgenic germline material was performed on embryos laid by the same mothers as for RNA seq, but at 15-16 days old, according to [63]. Small RNAs from strain 9 and strain 160 pooled ovaries were sequenced according to [64] with the oxidation reaction. Small RNAs from ovaries from individual F3 females was performed by Fasteris.

### Small RNA Analysis

Reads were trimmed by removing adapters and filtered by size as piRNA (23-30nt) in CLC Genomics Workbench 7.0. Reads were then filtered by mapping to tRNA, ncRNA, miscellaneous RNA, and miRNA (including pre-miRNA) libraries from the *D. virilis* reference genome. The filtered 23-30 nt small RNA reads were mapped to our curated TE library with BWA.aln[65], using the default parameters. Reads were normalized by non-unique mappers to the *D. virilis* reference genome using BWA.aln defaults. Calculations for ping-pong percent and density of piRNA pairs were done with the R package viRome (http://www.ark-genomics.org/bioinformatics/virome), with some modifications. For genic small RNA analysis, reads were mapped uniquely with BWA.aln to the *D. virilis* reference genome, using default parameters. Reads were normalized by non-unique mappers to the genome. BEDTools intersect[66] was utilized to count piRNA hits on genes and CDS sequences. Percent ping-pong was defined as the percent of 23 to 30 nt mapping reads that had a corresponding read on the opposite strand with a 10 bp 5’-5’ overlap. We also measured piRNA biogenesis by determining ping-pong pair density. This measure was obtained by counting all non-redundant ping-pong pairs (counting each read only once) per kb.

### Genetic analysis of genomic regions from strain 160 that maternally protect against dysgenesis

F3 females were generated by crossing 160 females to strain 9 males, followed by two rounds of backrossing to strain 9 males. All but the final cross was performed *en masse*. Dysgenic crosses were performed with >160 single 4 to 5 day old tester females mated with three 4 to 5 day old strain 160 males. Adults were transferred to new vials daily and dygenesis was estimated by counting the number of dysgenic testes in progeny over all testes counted (2 per male) across three broods. Females were then collected, ovaries removed (for small RNA sequencing by Fasteris, top protectors only) and carcasses retained for genomic DNA extraction. Genotyping was performed using the TaqMan Open Array platform on all females, with the exception of the top six females that had the strongest ability to protect against dysgenesis. SNPs distinguishing 160 and 9 chromosomes were chosen in pairs for redundancy, one pair at each telomere and pericentric region, as well as one or two euchromatic SNPs. Care was taken to avoid repeat sequences by screening with Blast against the reference and also using RepeatMasker. F3 Females were then genotyped for 160/9 heterozygosity, alongside pure strain 9 and 160 controls, by the National Jewish Health. Single marker regression was carried out with RQTL after dropping individuals with missing genotype data from the analysis. 5000 permutations were done to find the significance threshold at alpha = 0.05. For the top 6 protectors, whole genome sequencing was performed (100 bp, paired-end) using the Nextera library prep protocol.

### Genotyping by whole genome sequencing

Reference genome scaffolds from *D. virilis* were concatenated according to their supported positions and orientations on known Muller elements, with a large scaffold arbitrarily generated by concatenating scaffolds from unknown positions. Using this new “assembly” we mapped all strain 9 and strain 160 reads and generated two consensus genomes. A pseudo-diploid heterozygous genome was then assemble by placing these scaffolds into one file.

Paired-end reads from the top six protectors were mapped to the hybrid genome, using BWA’s default parameters, with the goal of inferring spans of heterozygosity for strain 160 by identifying reads that map uniquely, under high stringency, to the strain 160 haploid reference. The mapping output was piped into SAMtools for filtering by quality score (-q 42) and post-alignment processing (SAM to BAM conversion and indexing). A relatively high cutoff quality score was used in order to remove reads that could have mapped promiscuously. We were able to remove all reads that mapped to more than one locus/allele leaving us with reads that are specific to either strain 9 or 160. Spans of heterozygosity for strain 160 were visualized with a sliding window for read density along the 160 chromosomes within the psuedo-diploid reference genome.

## Acknowledgements

We thank Christine Yoder for early development of small RNA sequencing methods and Lena Hileman for providing comments on the manuscript. Jack Colicchio provided assistance in mRNA-seq analysis. Clark Bloomer from the KUMC Genomics core provided crucial assistance with DNA library prep and mRNA sequencing. We also thank Laura Rogers of the KU CMADP Genome Sequencing Core for additional mRNA sequencing. Additional DNA library prep, small RNA library prep and sequencing was performed by the David H. Murdock Research Institute and Fasteris. We also thank Daniel LaFlamme from National Jewish Health for assistance with genotyping. Funding was provided by NSF MCB-1022165, an internal grant through NIH-P20GM103638 and the University of Kansas.

## References

1. Hickey DA (1982) Selfish DNA: A sexually-transmitted nuclear parasite. Genetics 101: 519–531.

2. Charlesworth B, Charlesworth D (1983) The population dynamics of transposable elements. Genetical Research 42: 1–27.

3. Charlesworth B, Langley CH (1989) The population genetics of *Drosophila* transposable elements. Annual Review of Genetics 23: 251–287.

4. Kidwell MG, Novy JB (1979) Hybrid dysgenesis in *Drosophila melanogaster* - Sterility resulting from gonadal-dysgenesis in the P-M system. Genetics 92: 1127–1140.

5. Kidwell MG, Kidwell JF, Sved JA (1977) Hybrid Dysgenesis in *Drosophila melanogaster*: A Syndrome of Aberrant Traits Including Mutation, Sterility and Male Recombination. Genetics 86: 813–833.

6. Bingham PM, Kidwell MG, Rubin GM (1982) The molecular basis of P-M dysgenesis - The role of the P-element, a P-Strain-Specific Transposon Family. Cell 29: 995–1004.

7. Bucheton A, Paro R, Sang HM, Pelisson A, Finnegan DJ (1984) The molecular basis of the I-R hybrid dysgenesis syndreom in *Drosophila melanogaster -* Identification, cloning and properties of the I-Factor. Cell 38: 153–163.

8. Yannopoulos G, Stamatis N, Monastirioti M, Hatzopoulos P, Louis C (1987) *hobo* is responsible for the induction of hybrid dysgenesis by strains of *Drosophila melanogater* bearing the male recombination factor 23.5MRF. Cell 49: 487–495.

9. Engels WR (1989) P Elements in Drosophila melanogaster. In: Berg DE, Howe MH, editors. Mobile DNA. Washington, D.C.: American Society for Microbiology.

10. Lozovskaya ER, Scheinker VS, Evgenev MB (1990) A hybrid dysgenesis syndrome in *Drosophila virilis*. Genetics 126: 619–623.

11. Evgenev MB, Zelentsova H, Shostak N, Kozitsina M, Barskyi V, et al. (1997) *Penelope*, a new family of transposable elements and its possible role in hybrid dysgenesis in *Drosophila virilis*. Proceedings of the National Academy of Sciences of the United States of America 94: 196–201.

12. Pyatkov KI, Shostak NG, Zelentsova ES, Lyozin GT, Melekhin MI, et al. (2002) Penelope retroelements from Drosophila virilis are active after transformation of Drosophila melanogaster. Proceedings of the National Academy of Sciences of the United States of America 99: 16150–16155.

13. Evgen’ev MB (2013) What happens when Penelope comes?: An unusual retroelement invades a host species genome exploring different strategies. Mob Genet Elements 3: e24542.

14. Pyatkov KI, Arkhipova IR, Malkova NV, Finnegan DJ, Evgen’ev MB (2004) Reverse transcriptase and en:donuclease activities encoded by Penelope-like retroelements. Proceedings of the National Academy of Sciences of the United States of America 101: 14719–14724.

15. Petrov DA, Schutzman JL, Hartl DL, Lozovskaya ER (1995) Diverse transposable elements are mobilized in hybrid dysgenesis in *Drosophila virilis* Proceedings of the National Academy of Sciences of the United States of America 92: 8050–8054.

16. Vieira J, Vieira CP, Hartl DL, Lozovskaya ER (1998) Factors contributing to the hybrid dysgenesis syndrome in Drosophila virilis. Genetical Research 71: 109–117.

17. Aravin AA, Lagos-Quintana M, Yalcin A, Zavolan M, Marks D, et al. (2003) The small RNA profile during Drosophila melanogaster development. Developmental Cell 5: 337–350.

18. Grentzinger T, Armenise C, Brun C, Mugat B, Serrano V, et al. (2012) piRNA-mediated transgenerational inheritance of an acquired trait. Genome Res 22: 1877–1888.

19. Chambeyron S, Popkova A, Payen-Groschene G, Brun C, Laouini D, et al. (2008) piRNA-mediated nuclear accumulation of retrotransposon transcripts in the Drosophila female germline. Proc Natl Acad Sci U S A 105: 14964–14969.

20. Brennecke J, Malone CD, Aravin AA, Sachidanandam R, Stark A, et al. (2008) An Epigenetic Role for Maternally Inherited piRNAs in Transposon Silencing. Science 322: 1387–1392.

21. Khurana JS, Wang J, Xu J, Koppetsch BS, Thomson TC, et al. (2011) Adaptation to P Element Transposon Invasion in Drosophila melanogaster. Cell 147: 1551–1563.

22. Gerasimova T, Mizrokhi L, Georgiev G (1984) Transposition bursts in genetically unstable Drosophila melanogaster. Nature 309: 714–716.

23. Eggleston WB, Johnson-Schlitz DM, Engels WR (1988) P-M hybrid dysgenesis does not mobilize other transposable element families in D. melanogaster. Nature 331: 368–370.

24. Rozhkov NV, Zelentsova ES, Shostak NG, Evgen’ev MB (2011) Expression of Drosophila virilis retroelements and role of small RNAs in their intrastrain transposition. PLoS One 6: e21883.

25. Scheinker VS, Lozovskaya ER, Bishop JG, Corces VG, Evgenev MB (1990) A long terminal repeat-containing retrotransposon in mobilized during hybrid dysgenesis in *Drosophila virilis*. Proceedings of the National Academy of Sciences of the United States of America 87: 9615–9619.

26. Lyozin GT, Makarova KS, Velikodvorskaja VV, Zelentsova HS, Khechumian RR, et al. (2001) The structure and evolution of Penelope in the virilis species group of Drosophila: an ancient lineage of retroelements. Journal of Molecular Evolution 52: 445–456.

27. Blumenstiel JP (2014) Whole genome sequencing in Drosophila virilis identifies Polyphemus, a recently activated Tc1-like transposon with a possible role in hybrid dysgenesis. Mob DNA 5: 6.

28. Rozhkov NV, Schostak NG, Zelentsova ES, Yushenova IA, Zatsepina OG, et al. (2013) Evolution and dynamics of small RNA response to a retroelement invasion in Drosophila. Mol Biol Evol 30: 397–408.

29. Li H (2013) Aligning sequence reads, clone sequences and assembly contigs with BWA-MEM. arXivorg arXiv:1303.3997v2.

30. Daniels SB, Peterson KR, Strausbaugh LD, Kidwell MG, Chovnick A (1990) Evidence for horizontal transmission of the P transposable element between Drosophila species. Genetics 124: 339–355.

31. Morales-Hojas R, Vieira CP, Vieira J (2006) The evolutionary history of the transposable element Penelope in the Drosophila virilis group of species. J Mol Evol 63: 262–273.

32. Blumenstiel JP, Hartl DL (2005) Evidence for maternally transmitted small interfering RNA in the repression of transposition in Drosophila virilis. Proceedings of the National Academy of Sciences of the United States of America 102: 15965–15970.

33. Love MI, Huber W, Anders S (2014) Moderate estimation of fold change and dispersion for RNA-Seq data with DESeq2. bioRxiv.

34. Robinson MD, Oshlack A (2010) A scaling normalization method for differential expression analysis of RNA-seq data. Genome Biol 11: R25.

35. Robinson MD, McCarthy DJ, Smyth GK (2010) edgeR: a Bioconductor package for differential expression analysis of digital gene expression data. Bioinformatics 26: 139–140.

36. Eden E, Lipson D, Yogev S, Yakhini Z (2007) Discovering motifs in ranked lists of DNA sequences. PLoS Comput Biol 3: e39.

37. Eden E, Navon R, Steinfeld I, Lipson D, Yakhini Z (2009) GOrilla: a tool for discovery and visualization of enriched GO terms in ranked gene lists. BMC Bioinformatics 10: 48.

38. Kelleher ES, Edelman NB, Barbash DA (2012) Drosophila interspecific hybrids phenocopy piRNA-pathway mutants. PLoS Biol 10: e1001428.

39. Gu T, Elgin SC (2013) Maternal depletion of Piwi, a component of the RNAi system, impacts heterochromatin formation in Drosophila. PLoS Genet 9: e1003780.

40. Haynes KA, Caudy AA, Collins L, Elgin SCR (2006) Element 1360 and RNAi components contribute to HP1-dependent silencing of a pericentric reporter. Current Biology 16: 2222–2227.

41. Sentmanat MF, Elgin SC (2012) Ectopic assembly of heterochromatin in Drosophila melanogaster triggered by transposable elements. Proc Natl Acad Sci U S A 109: 14104–14109.

42. Sienski G, Donertas D, Brennecke J (2012) Transcriptional silencing of transposons by Piwi and maelstrom and its impact on chromatin state and gene expression. Cell 151: 964–980.

43. Olovnikov I, Ryazansky S, Shpiz S, Lavrov S, Abramov Y, et al. (2013) De novo piRNA cluster formation in the Drosophila germ line triggered by transgenes containing a transcribed transposon fragment. Nucleic Acids Res 41: 5757–5768.

44. Rozhkov NV, Aravin AA, Zelentsova ES, Schostak NG, Sachidanandam R, et al. (2010) Small RNA-based silencing strategies for transposons in the process of invading Drosophila species. Rna-a Publication of the Rna Society 16: 1634–1645.

45. Ronsseray S, Josse T, Boivin A, Anxolabehere D (2003) Telomeric transgenes and trans-silencing in Drosophila. Genetica 117: 327–335.

46. Ronsseray S, Marin L, Lehmann M, Anxolabehere D (1998) Repression of hybrid dysgenesis in Drosophila melanogaster by combinations of telomeric P-element reporters and naturally occurring P elements. Genetics 149: 1857–1866.

47. Marin L, Lehmann M, Nouaud D, Izaabel H, Anxolabehere D, et al. (2000) P-element repression in Drosophila melanogaster by a naturally occurring defective telomeric P copy. Genetics 155: 1841–1854.

48. Simmons MJ, Ragatz LM, Sinclair IR, Thorp MW, Buschette JT, et al. (2012) Maternal enhancement of cytotype regulation in Drosophila melanogaster by genetic interactions between telomeric P elements and non-telomeric transgenic P elements. Genet Res (Camb) 94: 339–351.

49. Niemi JB, Raymond JD, Patrek R, Simmons MJ (2004) Establishment and maintenance of the P cytotype associated with telomeric P elements in Drosophila melanogaster. Genetics 166: 255–264.

50. Rozhkov NV, Aravin AA, Zelentsova ES, Schostak NG, Sachidanandam R, et al. (2010) Small RNA-based silencing strategies for transposons in the process of invading Drosophila species. RNA 16: 1634–1645.

51. Brennecke J, Aravin AA, Stark A, Dus M, Kellis M, et al. (2007) Discrete small RNA-generating loci as master regulators of transposon activity in Drosophila. Cell 128: 1089–1103.

52. de Vanssay A, Bouge AL, Boivin A, Hermant C, Teysset L, et al. (2012) Paramutation in Drosophila linked to emergence of a piRNA-producing locus. Nature 490: 112–115.

53. Arteaga-Vazquez M, Sidorenko L, Rabanal FA, Shrivistava R, Nobuta K, et al. (2010) RNA-mediated trans-communication can establish paramutation at the b1 locus in maize. Proc Natl Acad Sci U S A 107: 12986–12991.

54. Alleman M, Sidorenko L, McGinnis K, Seshadri V, Dorweiler JE, et al. (2006) An RNA-dependent RNA polymerase is required for paramutation in maize. Nature 442: 295–298.

55. Chandler VL (2007) Paramutation: from maize to mice. Cell 128: 641–645.

56. Mott R, Yuan W, Kaisaki P, Gan X, Cleak J, et al. (2014) The architecture of parent-of-origin effects in mice. Cell 156: 332–342.

57. Gibson G, Riley-Berger R, Harshman L, Kopp A, Vacha S, et al. (2004) Extensive sex-specific nonadditivity of gene expression in Drosophila melanogaster. Genetics 167: 1791–1799.

58. Wittkopp PJ, Haerum BK, Clark AG (2006) Parent-of-origin effects on mRNA expression in Drosophila melanogaster not caused by genomic imprinting. Genetics 173: 1817–1821.

59. Engels WR (1979) Hybrid dysgenesis in *Drosophila melanogaster* - Rules of inheritance of female sterility. Genetical Research 33: 219–236.

60. Smith CD, Edgar RC, Yandell MD, Smith DR, Celniker SE, et al. (2007) Improved repeat identification and masking in Dipterans. Gene 389: 1–9.

61. Li R, Ye J, Li S, Wang J, Han Y, et al. (2005) ReAS: Recovery of ancestral sequences for transposable elements from the unassembled reads of a whole genome shotgun. PLoS Comput Biol 1: e43.

62. Clark AG, Eisen MB, Smith DR, Bergman CM, Oliver B, et al. (2007) Evolution of genes and genomes on the *Drosophila* phylogeny. Nature 450: 203–218.

63. Vigneault F, Ter-Ovanesyan D, Alon S, Eminaga S, D CC, et al. (2012) High-throughput multiplex sequencing of miRNA. Curr Protoc Hum Genet Chapter 11: Unit 11 12 11-10.

64. Li CJ, Vagin VV, Lee SH, Xu J, Ma SM, et al. (2009) Collapse of Germline piRNAs in the Absence of Argonaute3 Reveals Somatic piRNAs in Flies. Cell 137: 509–521.

65. Li H, Durbin R (2010) Fast and accurate long-read alignment with Burrows-Wheeler transform. Bioinformatics 26: 589–595.

66. Quinlan AR, Hall IM BEDTools: a flexible suite of utilities for comparing genomic features. Bioinformatics 26: 841–842.

